# The Resolvin D2-GPR18 Axis Enhances Bone Marrow Function and Limits Hepatic Fibrosis in Aging

**DOI:** 10.1101/2023.01.05.522881

**Authors:** Hannah Fitzgerald, Jesse L. Bonin, Sudeshna Sadhu, Masharh Lipscomb, Nirupam Biswas, Christa Decker, Melisande Nabage, Ramon Bossardi, Michael Marinello, Agustina Hebe Mena, Kurrim Gilliard, Matthew Spite, Alejandro Adam, Katherine C. MacNamara, Gabrielle Fredman

**Affiliations:** The Department of Molecular and Cellular Physiology, Albany Medical College, Albany, NY 12208, USA; The Department of Immunology and Microbial Disease, Albany Medical College, Albany, NY 12208, USA; Center for Experimental Therapeutics and Reperfusion Injury, Department of Anesthesiology, Perioperative and Pain Medicine, Brigham and Women’s Hospital and Harvard Medical School, Boston, MA 02115, USA

**Keywords:** resolvin, inflammation, resolution, liver, monocyte, macrophage, bone marrow

## Abstract

Aging is associated with non-resolving inflammation and tissue dysfunction. Resolvin D2 (RvD2) is a pro-resolving ligand that acts through the G-protein coupled receptor (GPCR) called GRP18. Using an unbiased screen, we report increased *Gpr18* expression in macrophages from old mice and in livers from elderly humans that is associated with increased steatosis and fibrosis in middle-aged (MA) and old mice. MA mice that lack GPR18 on myeloid cells had exacerbated steatosis and hepatic fibrosis, which was associated with a decline in Mac2+ macrophages. Treatment of MA mice with RvD2 reduced steatosis and decreased hepatic fibrosis, correlating with increased Mac2+ macrophages, monocyte-derived macrophages and elevated numbers of monocytes in the liver, blood, and bone marrow. RvD2 acted directly upon the bone marrow to increase monocyte-macrophage progenitors. Using a transplantation assay we further demonstrated that bone marrow from old mice facilitated hepatic collagen accumulation in young mice, and transient RvD2 treatment to mice transplanted with bone marrow from old mice prevented hepatic collagen accumulation. Together, our study demonstrates that RvD2-GPR18 signaling controls steatosis and fibrosis and provides a mechanistic-based therapy for promoting liver repair in aging.

## Introduction

Aging is associated with chronic, non-resolving inflammation that can lead to tissue dysfunction(1, 2). Therefore, understanding cellular programs and factors that promote the resolution of inflammation during aging may inform the development of new strategies that can limit age-related organ decline. The resolution of inflammation is an active process that is governed by numerous factors, such as specialized pro-resolving lipid mediators (SPMs)(3). Recent results suggest that non-resolving inflammation in aging, or “inflammaging”, may persist due to an impairment in inflammation-resolution programs and that treatment with SPMs like Resolvins temper exuberant inflammation and age-related tissue dysfunction (2, 4, 5). However, little is known about SPM-initiated mechanisms that limit features of inflammaging.

SPMs, like Resolvin D2 (RvD2), promote a pro-resolving and pro-reparative phenotype in human macrophages and in young mice in the context of infection and injury (6-10). RvD2 acts through its G-protein coupled receptor (GPCR) called GPR18, which is expressed on macrophages in both humans and mice (8). RvD2 regulates phagocytosis, adhesion receptor expression, and cytokine production in phagocytes (6-8). The RvD2-GPR18 signaling axis is not immunosuppressive and boosts host immunity in polymicrobial sepsis (7) and in *E. coli* and *Staphylococcus aureus* (6, 8) infections in mice and promotes tissue repair and regeneration in young mice (10). Thus, the RvD2-GPR18 axis tames inflammation, does not compromise host defense, and promotes tissue repair in specific contexts, yet how this pathway acts in aging is not known.

Inflammaging is associated with chronic activation of the innate immune system characterized by elevated levels of circulating pro-inflammatory cytokines (1, 11). Despite heightened innate immune programs, elderly individuals have reduced host defenses, limited vaccine efficacy, an overall decline of tissue function and poor organ transplant efficacy (12, 13). Therefore, understanding mechanisms to limit tissue decline and/or boost tissue repair without comprising host defense in the elderly is an unmet clinical need. Monocytes and macrophages are considered integral cell types in the etiology of inflammaging (14). The aging milieu typically primes monocytes and macrophages towards a pro-inflammatory phenotype and, as such, macrophages in this context participate in propagating inflammation by secreting pro-inflammatory cytokines and lipid mediators (15). How to reprogram monocytes and macrophages toward a more pro-resolving and pro-reparative phenotype is of considerable interest in aging.

A consequence of inflammaging is organ dysfunction. The liver, for example, is one of the largest solid organs in the body that regulates metabolism and immunity(16) and an organ where age significantly increases the risk of fibrosis (17). Macrophages are the most abundant immune cell in the liver where they play critical roles in inflammation, fibrosis or fibrosis resolution (18, 19). Aging is associated with maladaptive structural and functional modifications in the liver. For example, blood volume in the liver decreases and the exchange of molecules becomes hampered due to the capillarization of liver sinusoids. Livers from elderly individuals are also associated with increased steatosis and collagen (fibrosis). Indeed, non-alcoholic fatty liver diseases (NAFLD) and nonalcoholic steatohepatitis (NASH) mainly affect the middle-aged and elderly populations (20, 21). Therefore, understanding how to limit age-related hepatic fibrosis is of clinical interest.

Here, using an unbiased screen, we report increased *Gpr18* expression in macrophages from old mice and in livers from elderly humans that is associated with increased steatosis and fibrosis in middle-aged (MA) and old mice. MA mice that lack GPR18 on myeloid cells had exacerbated steatosis and hepatic collagen accumulation, which was associated with a decline in monocytes. Treatment of MA mice with RvD2 reduced liver fat and collagen accumulation, correlating with increased monocyte-derived macrophages and elevated numbers of monocytes in the liver, blood, and bone marrow. RvD2 acted directly upon the bone marrow to increase monocyte-macrophage progenitors. Using a transplantation assay we demonstrated that bone marrow from old mice induced collagen accumulation in the liver of young mice, and transient RvD2 treatment limited the ability of marrow to drive this aging phenotype. Together, our study demonstrates that RvD2-GPR18 signaling controls age-related steatosis and fibrosis and provides a mechanistic-based therapy for promoting liver repair in aging.

## Results

### Gpr18 expression is increased in macrophages from old mice and in livers from old humans

Macrophages are critical cellular players of both inflammation and resolution, so we questioned whether macrophages from old mice had distinct gene expression from young mice. Zymosan-elicited macrophages were harvested from young (2 month) and old (18 month) mice and subjected to bulk RNA sequencing. Principal component analysis (PCA) revealed that peritoneal macrophages elicited from old mice have distinct gene signatures compared with peritoneal macrophages from young mice (**Fig. 1A**). Gene set enrichment analysis (GSEA) revealed upregulated pathways associated with inflammation (e.g. COX2), MAPK signaling (e.g. p38) and ROS (**Supp Fig. 1**), which is consistent with other observations of age-related changes to macrophages (19). GPR18 (i.e. the receptor for RvD2) was among the genes that were significantly upregulated (**Fig. 1B**). Additionally, bone marrow-resident macrophages from old mice (22) had significantly increased *Gpr18* expression compared with those from young mice (**Fig. 1C**). To test whether macrophages from old mice were responsive to RvD2, young or old mice were intraperitoneally injected with RvD2 (250 ng/mouse/day for 7 days) and peritoneal macrophages from young and old mice were collected by lavage. First, we found that macrophages from old mice had increased levels of COX-2 (**Supp Fig. 1B**), phospho-p38 (**Supp Fig. 1C**) and H2AXy (**Supp Fig. 1D**) compared with macrophages from young mice, and RvD2 significantly decreased COX-2, phospho-p38, and H2AXy. These data demonstrate that macrophages from old mice have increased *Gpr18* expression and are responsive to RvD2 stimulation, suggesting that the RvD2-GPR18 axis, specifically in macrophages, may be a targetable signaling pathway to limit inflammaging.

**Fig 1.**
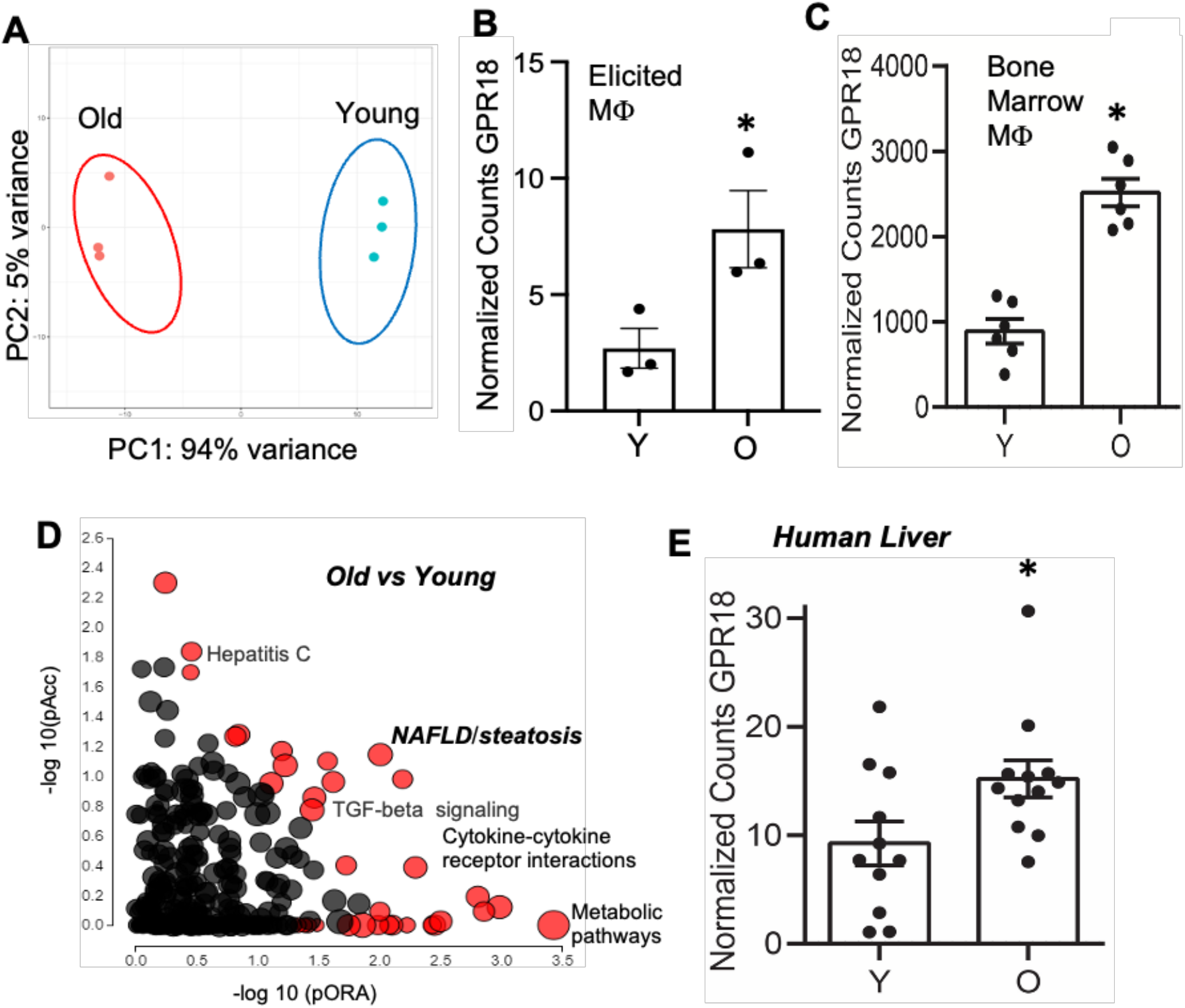
Differentially expressed genes and pathways in zymosan-elicited peritoneal macrophages from young and old mice. RNA sequencing was performed on Zymosan-elicited peritoneal macrophages from young (2 mo) or old (18 mo) mice as described in the methods. (**A**) Principal Component Analysis (PCA) for the young or aged mice is shown. (**B**,**C**) Normalized counts for *Gpr18* expression in (**B**) Zym-elicited macrophages and (**C**) murine bone marrow macrophages are shown. Each dot represents an individual mouse and *p ≤ 0.05, Student’s *t test*. (**D**) iPathwayGuide analysis shows biological pathways with significant perturbation (pAcc) and overrepresentation (pORA) in old as compared to young mice. Significant pathways are shown in red and non-significant pathways are shown in black. (**E**) Normalized counts of GPR18 expression in human liver. LogFC 6.12, p-value = 0.03.

Additionally, differential gene expression analysis (iPathway Guide) suggested that several of the significantly upregulated pathways in macrophages from aged mice were associated with inflammation, metabolism, and liver pathology, such as non-alcoholic fatty liver disease (NAFLD) (**Fig. 1D**). Human livers from young (21-45 yrs.) and old (69+ yrs.) individuals were examined for *GPR18* expression and we found that livers from old individuals had significantly increased expression of *GPR18* (**Fig. 1E**). Collectively, these data suggest that elevated *GPR18* levels in macrophages from old mice and livers from old humans may yield important clues about new targetable approaches toward limiting inflammaging. Moreover, these data focused our questions as to how macrophages and GPR18 signaling may play a role in age-related liver pathology.

### Middle-aged mice have increased liver collagen and fatty deposits

Based on the results from our unbiased screen, indicating an upregulation in pathways associated with NAFLD, we next thought to assess lipid content in the liver from young (Y, 2-3 month), middle-aged (MA, 11-12 month), and old (O, 17-19 month) mice fed a standard chow diet. Of note, body weight significantly increased with age as expected (Y: 18.6g ± 1.7; MA: 32.7g ± 3.2; O: 42.3g ± 3.6). We quantified lipid accumulation in the liver with Oil Red O staining (shown as red) and image analysis revealed a statistically significant increase in Oil Red O staining in MA and old livers compared with young livers (**Fig. 2A**). H&E staining revealed increased fatty deposits in MA and old livers (**Fig. 2B, left**). To assess liver triglycerides (TGs), livers from each group were homogenized and assayed for TG levels as described in the methods. MA and old mice had significantly increased liver TGs compared with young mice (**Fig. 2B, right**). These data demonstrate that accumulation of fat in liver occurs as early as middle age.

**Fig. 2.**
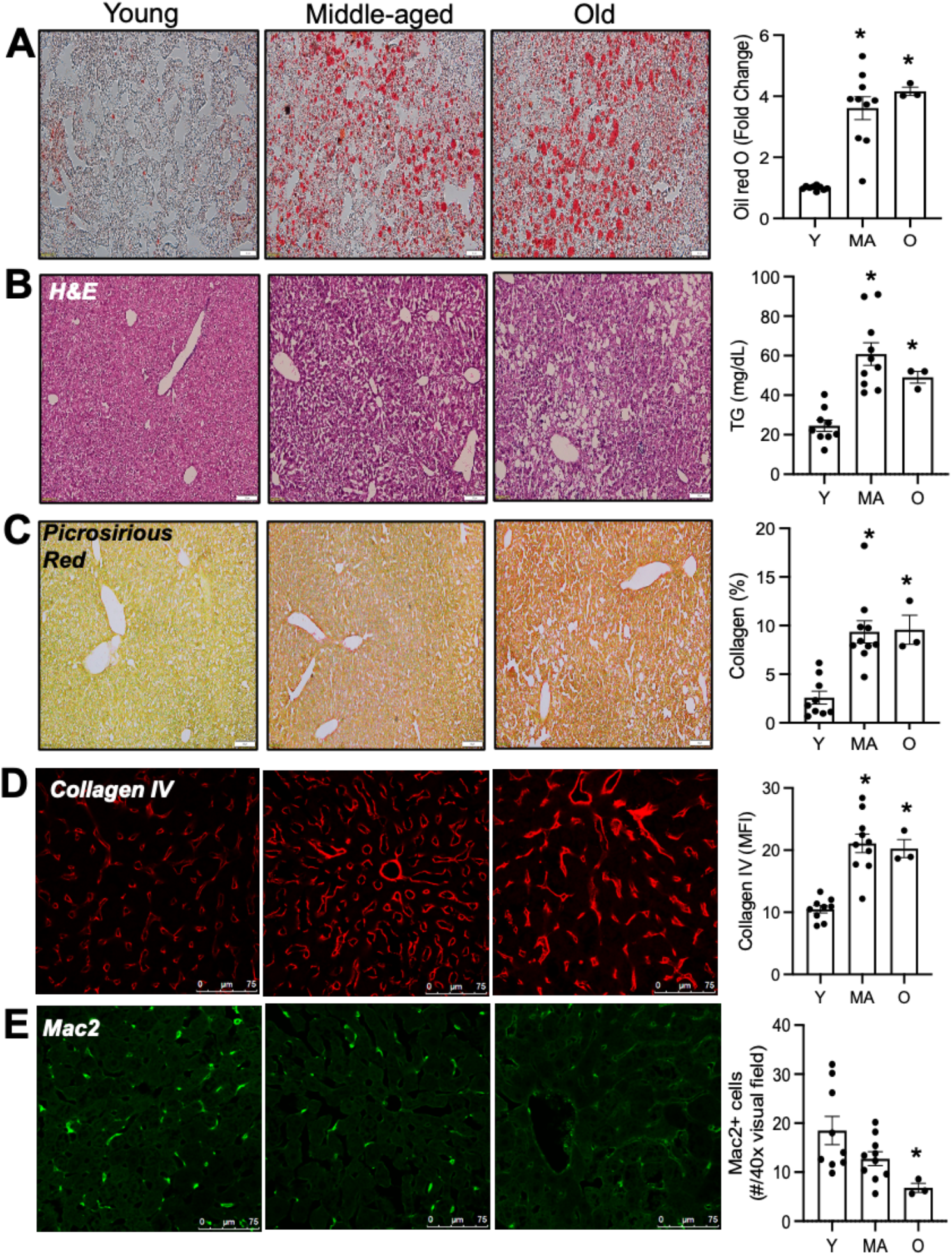
Middle-aged and old mice have elevated steatosis and hepatic fibrosis compared with young mice. Livers from young (Y, 2 months), middle-aged (MA, 11 months) or old (O, 18 months) mice were harvested (**A**) Frozen liver sections were stained with Oil Red O and lipid deposits are visualized as red. Scale bar is 20μm. Images were acquired on a Olympus microscope and red content was analyzed with Image J. (**B, left**) Liver sections were stained with H&E and representative images are shown. Scale bar is 100μm. (**B, right**) Liver homogenates were assessed for triglyceride content. (**C**) Representative images and quantification of picrosirius red. Scale bar is 100μm. Images were acquired on an Olympus microscope and the quantification was done by Image J analysis and are presented as the % of red area per 4x visual field (**D**) Representative images and quantification of liver sections that were immunostained with anti-Collagen IV (Col-IV) antibody (red) or (**E**) anti-Mac2 antibody (green). (**D-E**) Images were acquired on a Leica confocal microscope and quantification of (**D**) Col-IV mean fluorescence intensity (MFI) and (E) the number of Mac2+ cells/40x visual field were enumerated with Image J software. The scale bar for (**D** and **E)** is 75μm. (Results are mean ± SEM and each dot represents an individual mouse. *p < 0.05, Kruskal-Wallis and Dunn’s post-test.

Another histological feature of liver aging, as with other key organs, is fibrosis. Livers were stained with picrosirius red for the identification of collagen networks and images depict increased red color which was significantly increased in the livers from MA and O mice (**Fig. 2C**) compared with Y mice. Indeed, liver fibrosis is associated with accumulation of collagens I, III and IV (23) and collagen IV is a fibrillar collagen that is a major extracellular matrix (ECM) component of the basement membrane. Liver sinusoids are unique because they are lined by a fenestrated endothelium that lack a basement membrane, and formation of perisinusoidal basement membranes beneath the endothelium is an integral feature of capillarization of sinusoids that is associated with increased fibrosis in livers from elderly individuals (24). Therefore, we next used an immunofluorescence-based approach where livers were immunostained with an anti-collagen-IV antibody. Representative images revealed increased Col-IV signal (as shown in red) and quantification demonstrated a significant increase in Col-IV staining in MA and old mice compared with young mice (**Fig. 2D**). Together, these data suggest that hepatic fibrosis occurs in mice as early as middle age.

### Middle-aged mice have decreased liver Mac2+ macrophages

Macrophages can promote fibrogenesis and resolve fibrosis (18) and so we questioned whether aging was associated with differences in macrophages in the liver. Using immunofluorescence we found that Mac2 (Galectin-3, or Gal3) staining, which identifies macrophages, decreased with age (**Fig. 2E**). While we noted a trend for decreased Mac2^+^ cells in MA mice, the decrease reached statistical significance in old mice (**Fig. 2E**). Mac2-expressing macrophages are phagocytic (25) and therefore the loss of Mac2 expression in livers from aging mice may be indicative of a decline in phagocytic macrophages. The absolute frequency of liver macrophages, as measured by F4/80 staining of total liver cells and flow cytometric analysis, did not change with age (Y: 7.9% ± 1.9; MA: 8.2% ± 1.8; O: 9% ± 2.9), suggesting that the population of Mac2+ macrophages decreased with age. Together, these data suggest an association between decreased Mac2^+^ macrophages and increased steatosis and hepatic fibrosis in middle-age livers compared with young controls.

### Myeloid Gpr18 limits fat and collagen accumulation in livers of aging mice

Because aging is associated with impaired inflammation-resolution programs (2, 4), and because we found that *Gpr18* expression was increased in macrophages from old mice and in livers from elderly humans, we next questioned whether the above changes we observed in mice during the aging process were due to impairments in GPR18 singling. To evaluate the role of GPR18 in myeloid cells in age-associated liver pathology, we generated a novel mouse model wherein a floxed human *GPR18* allele was inserted to the mouse genome, replacing the murine *Gpr18* gene (**Supp. Fig. 2A**). The human floxed *GPR18* gene (referred to as hGPR18fl/fl or “fl/fl”) was crossed with a *LysM*-driven Cre line to generate myeloid-specific GPR18 knockout mice (hGPR18mKO or “mKO”). First, we generated bone marrow-derived macrophages (BMDMs) from wild type C57BL/6, fl/fl and mKO mice to determine expression levels of both murine and human *GPR18*. We confirmed a significant loss of murine *Gpr18* by qPCR in BMDMs from both fl/fl and mKO mice (**Supp Fig. 2B**), and human *GPR18* was primarily expressed in the fl/fl mice (**Supp Fig. 2C**). To determine whether human GPR18 was functional in murine cells, we next performed an *in vitro* phagocytosis assay. Briefly, BMDMs were treated with Veh or RvD2 (1 nM) for 15 mins, after which opsonized fluorescent zymosan particles were added. After 30 mins, the fluorescent zymosan particles were washed off and phagocytosis was assessed by a fluorescence plate reader. We found that RvD2 enhanced phagocytosis in BMDMs from C57BL6 control mice and the fl/fl mice, suggesting that the human GPR18 was as functional as murine GPR18 in murine cells (**Supp Fig. 2D**). Importantly, RvD2 did not increase phagocytosis in the mKO BMDMs, demonstrating that we successfully removed both murine and human GPR18 in macrophages and that RvD2’s ability to enhance phagocytosis was dependent on GPR18.

We next asked whether GPR18 signaling on myeloid cells impacts steatosis and hepatic fibrosis in MA mice. Female fl/fl and mKO mice were aged until 10-12 months and their livers assessed for TG, collagen, Mac2+ macrophages and monocytes as described above. First, MA mKO had had significantly more Oil Red O staining compared with fl/fl middle-aged controls (**Fig. 3A**), despite no changes in body weight between the groups (fl/fl 33.8g ± 5.2 vs mKO 35.9g ± 4.0 p = 0.57). MA mKO mice also had a significant increase in Col-IV staining relative to the fl/fl controls (**Fig. 3B**). Together, these data suggest that the loss of GPR18 on myeloid cells drives steatosis and hepatic collagen accumulation in middle-aged mice.

**Fig. 3.**
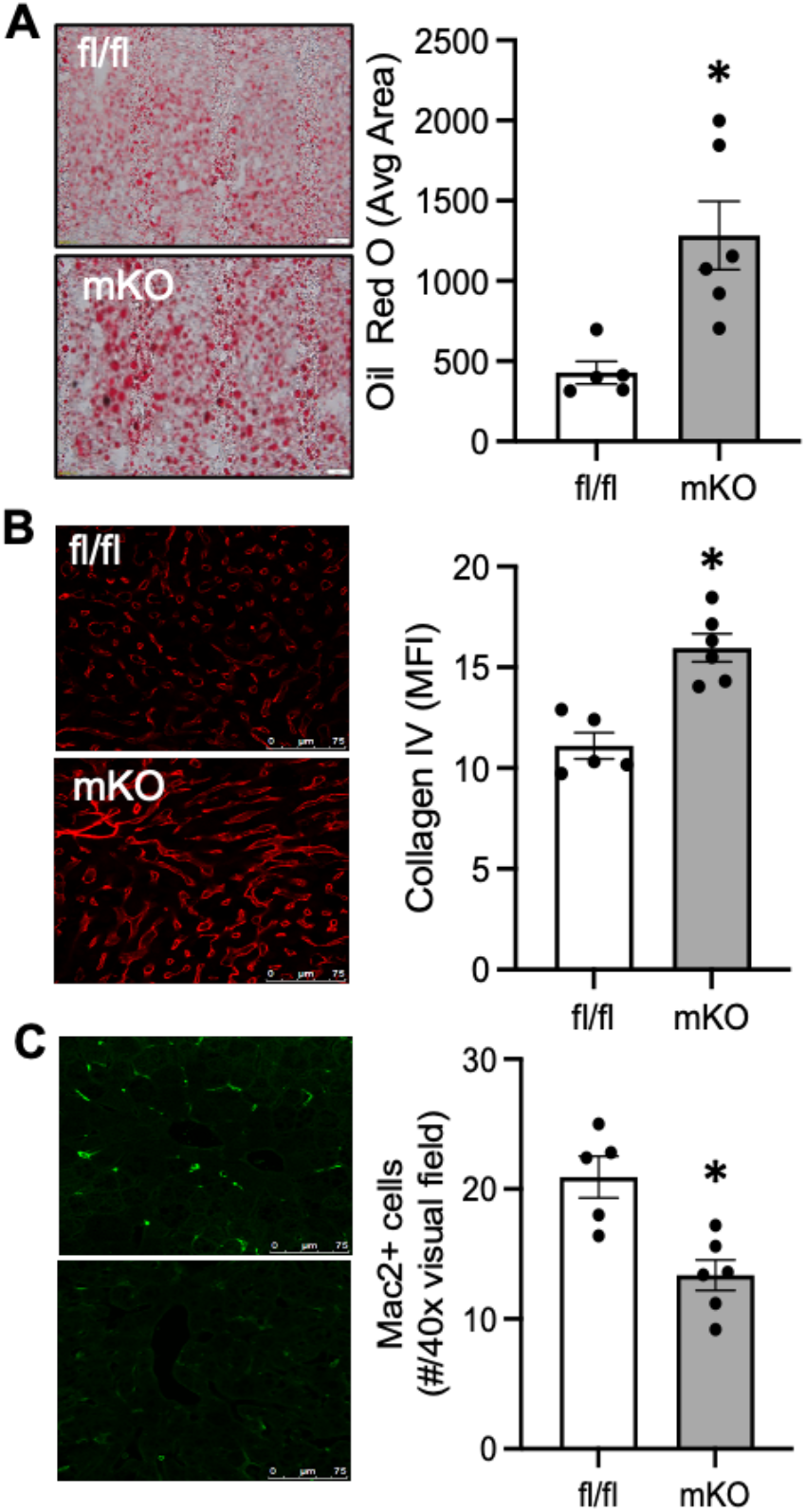
Middle-aged *Gpr18* mKO mice have increased liver triglyceride and collagen and decreased liver monocytes compared with middle-aged fl/fl mice. (**A**) Frozen liver sections were stained with Oil Red O and lipid deposits are visualized as red. Scale bar is 20μm. Oil Red O was quantified as in Fig. 2. Livers from MA fl/fl and mKO mice were immunostained for (**B**) Col-IV and (**D**) Mac2. Scale bar is 75μm. Images were acquired on a Leica confocal microscope and quantification of Col-IV MFI or Mac2+ positive cells was done with Image J software. Results are mean ± SEM and each dot represents an individual mouse. *p< 0.05 Mann-Whitney.

Because we found a decreasing trend for Mac2^+^ macrophages in MA, we next questioned whether the loss of GPR18 on myeloid cells would impact these cells in the liver. Liver sections were stained with Mac2 as above and indeed, MA mKO livers had significantly fewer Mac2^+^ cells, compared with fl/fl MA controls (**Fig. 3C**). Collectively, these data reveal a key role for cell-autonomous RvD2-GPR18 signaling in myeloid cells in controlling age-related steatosis and hepatic fibrosis. Thus, we reasoned that targeting GPR18 could offer a novel therapeutic strategy.

### Resolvin D2 limits age-related steatosis and collagen accumulation in the liver

To address the possibility of targeting GPR18 therapeutically, we tested whether RvD2 treatment impacted the liver in MA mice. MA mice were given Veh or RvD2 (250 ng/mouse, i.p.) for 7 days. Mice were then sacrificed and tissues were collected for analysis. Livers were sectioned and stained with H&E, which revealed reduced appearance of fat deposits in RvD2-treated MA mice (**Fig. 4A**) and significantly decreased liver TG levels (**Fig. 4B**), without a change in body weight (**Supp Fig. 3A**). Picrosirious red-staining revealed reduced collagen deposition in the perivascular regions, and quantification revealed a statistically significant decrease in picrosirius red staining in MA + RvD2 compared mice with MA vehicle controls (**Fig. 4C**). Immunofluorescence of livers for Col-IV, as shown in red in the representative images, was also significantly decreased by RvD2 (**Fig. 4D**). RvD2 also significantly decreased circulating C-reactive protein (CRP) levels in MA mice compared with young mice (**Supp Fig. 3B**). Together, these data demonstrate a significant improvement in age-related signatures of steatosis hepatic fibrosis and inflammation with one week of RvD2 treatment.

**Fig. 4.**
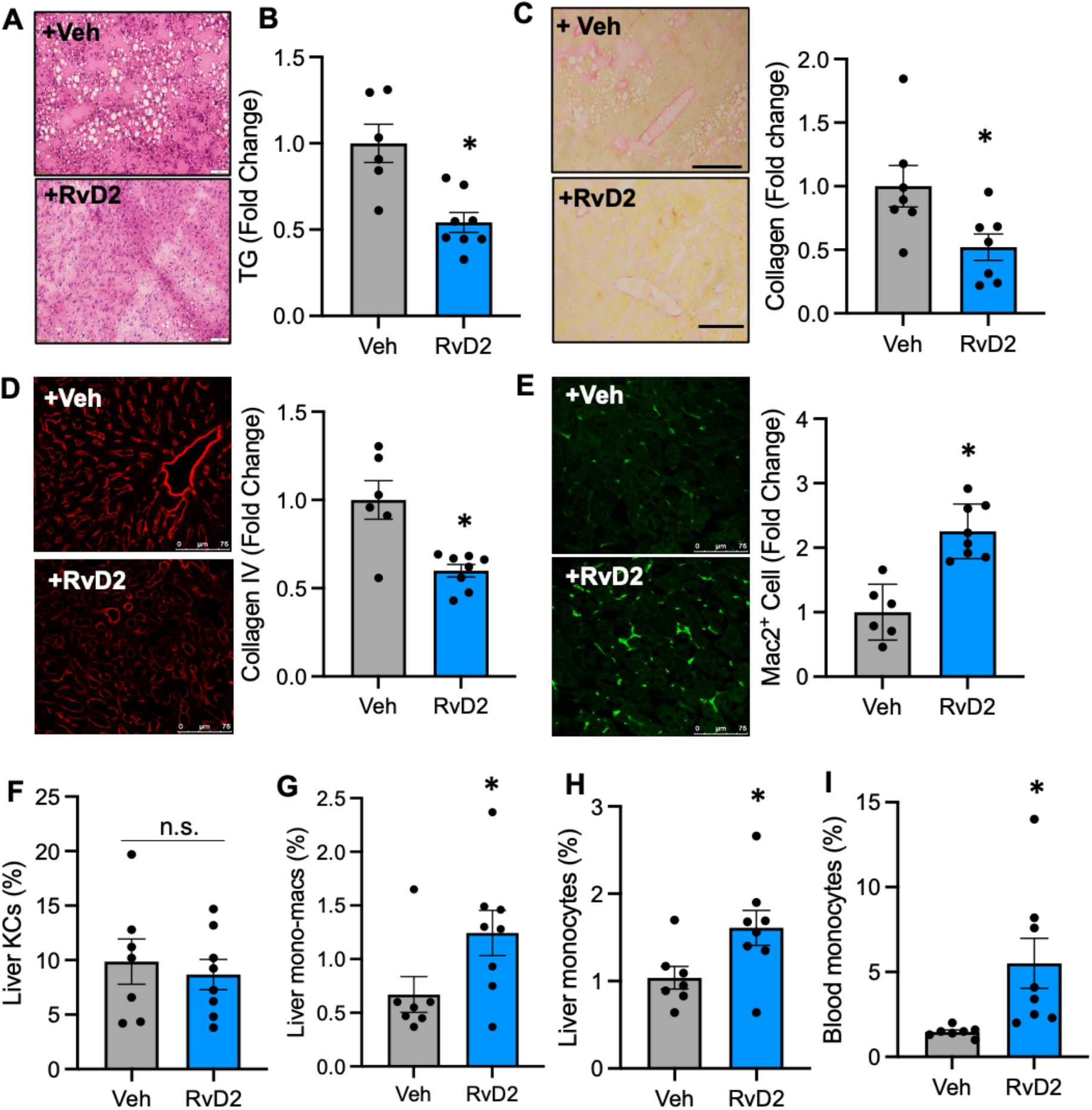
RvD2 limits steatosis, reduces hepatic fibrosis and increases Mac2+ macrophages and monocytes in the livers from middle-aged mice. MA (11mo) mice were treated with Veh or RvD2 (250ng/mouse, i.p.) for 7 days. Livers were harvested as in Fig 2. (**A**) Liver sections were stained with H&E are representative images are shown. Scale bar is 100μm. (**B**) Liver homogenates were assessed for triglyceride content. (**C**) Picrosirious red was analyzed by Image J analysis and are represented as fold change of Veh treated mice. Scale bar is 200μm. (**D**) Col-IV mean fluorescence intensity (MFI) and (**E**) Mac2+ cells were analyzed with Image J software and represented as fold change of Veh. (**F-H**) Liver was processed and analyzed by flow cytometry for (**F**) Kupffer cells (Ly6G^-^, CD11b^lo^, F4/80^hi^), (**G**) monocyte-derived macrophages (Ly6G^-^, CD11b^+^, F4/80^+^) and (**H**) monocytes (Ly6G^-^, CD11b^+^, Ly6C^hi^). (**I**) Circulating monocytes were quantified by a Hesta-HT5 analyzer. Results are mean ± SEM and each dot represents an individual mouse. *p< 0.05 Mann-Whitney.

Consistent with a potential role for macrophages in age-induced liver pathology, we found that RvD2-treated MA mice had significantly increased Mac2^+^ cells in the liver as determined by immunofluorescence and confocal imaging (**Fig 4E**). By flow cytometry (**Supp. Fig. 4**) we observed that resident KCs were not changed between the two groups (**Fig. 4F**), however, RvD2 significantly increased monocyte-derived macrophages (CD11b^hi^, F4/80+) (**Fig. 4G**) and liver monocytes (**Fig. 4H**). We next questioned whether the increased monocytes in the liver were due to systemic increases in circulating monocytes. Complete blood count (CBC) analysis revealed that RvD2 treatment significantly increased the frequency of monocytes among circulating white blood cells (WBCs) in the blood (**Fig. 4I**). Collectively, these data suggest that increased liver Mac2+ macrophages, monocyte-derived macrophages and monocytes in RvD2-treated mice correlates with decreases steatosis and hepatic fibrosis in MA mice.

### RvD2 increased monocytes in bone marrow and promotes monopoiesis

Monocytes are found in circulation where they can be recruited into tissues to participate in replacement of macrophages. Monocytes were increased not only in livers from RvD2-treated MA mice, but also in the blood. Therefore, we next questioned whether monocytes were also increased in the bone marrow where they are produced. Middle-aged mice treated with RvD2 exhibited a significant increase frequency (**Fig. 5A**) and number of bone marrow monocytes (**Fig. 5B**). To determine if RvD2 impacted hematopoietic progenitors we analyzed phenotypic hematopoietic stem and progenitor cells (HSPCs) by flow cytometry in MA mice with or without RvD2 treatment (**Supp. Fig 5**). RvD2 induced an increase in the pool of Lineage-negative cKit+ Sca1+ (LSK) cells, however within the LSK population we found that RvD2 reduced the frequencies of phenotypic hematopoietic stem cells (HSCs) and multipotent progenitors (MPPs) (**Fig. 5C**). At the same time, RvD2 increased phenotypic, myeloid-biased MPPs (MPP_GM_) within the LSK population (**Fig. 5C**). Moreover, RvD2 significantly increased absolute frequencies (**Fig. 5D**) and numbers of myeloid-committed MPP_GM_ cells (**Fig. 5E**) in the marrow. These data suggest that RvD2 treatment of MA mice induced an expansion of myeloid lineage-committed progenitors.

**Fig. 5.**
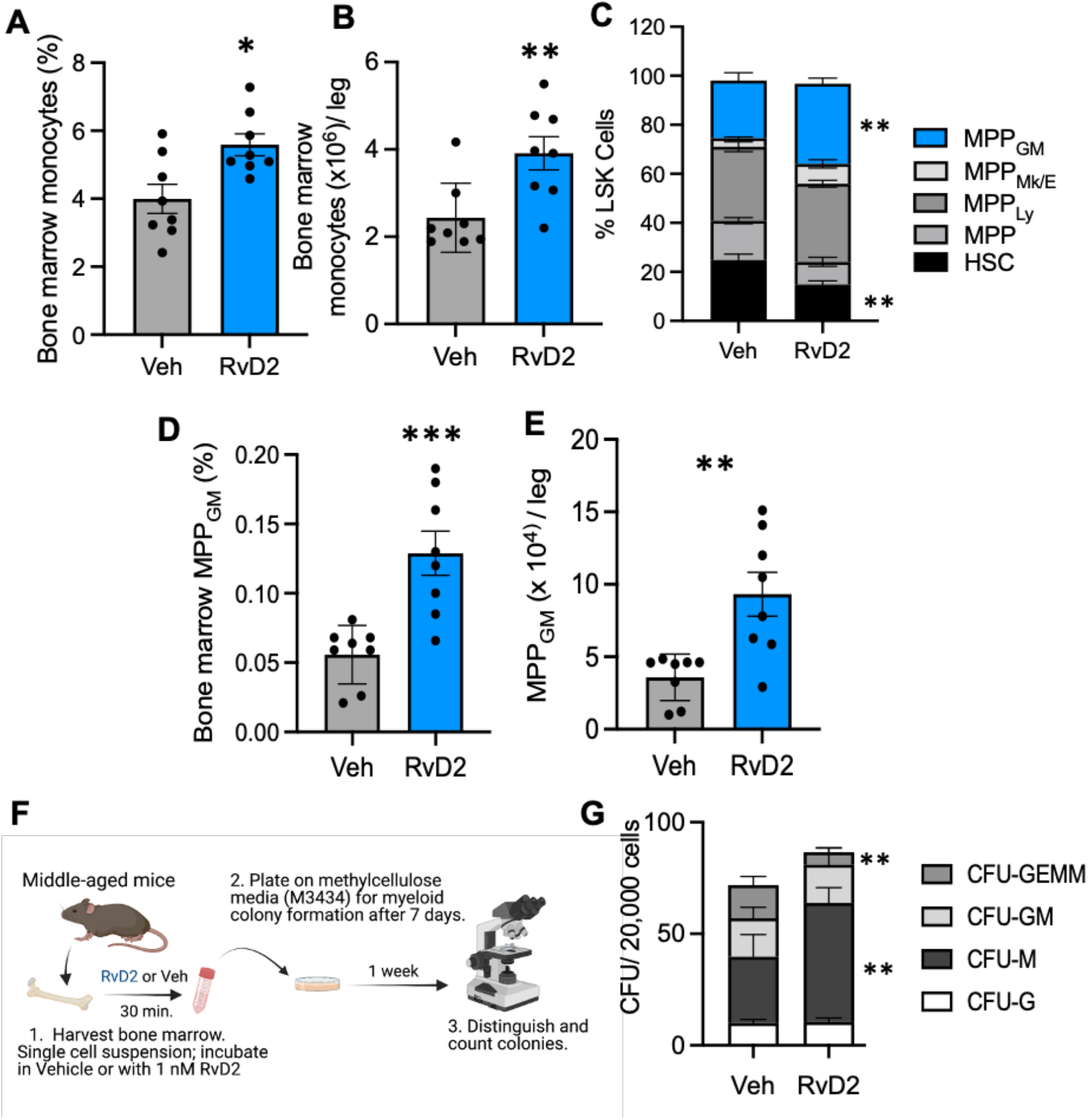
RvD2 Increases bone marrow monocytes and monocyte progenitors. Bone marrow was harvested and monocytes were enumerated by flow cytometry. (**A-B**) Monocyte (CD11b^+^ Ly6G^-^ Ly6C^hi^) frequencies and numbers in MA mice treated with vehicle or RvD2 are shown. (**C**) Frequencies of phenotypic HSPCs in bone marrow are shown as the percent of each population among total Lineage-negative, c-Kit+, and Sca-1+ cells (LSK). (**D**,**E**) The absolute frequencies (**D**) and number (**E**) of granulocyte-macrophage multipotent progenitors (MPP_GM_) are shown. (**F**) Bone marrow was obtained from MA mice and incubated for 30 minutes with vehicle or RvD2 (1 nM) and plated in methocult media. After 7 days colonies were counted. (**G**) CFUs are shown for CFU-M, CFU-G, CFU-GM, and CFU-GEMM from middle-aged bone marrow treated with vehicle or RvD2. Results are mean ± SEM and each dot represents an individual mouse. *p< 0.05, **p< 0.01 Mann-Whitney. For experiments conducted in (**G**), Results are mean ± SEM of bone marrow from n = 4 separate mice per treatment.

To examine the potential impact of RvD2 on functional myelopoiesis we isolated bone marrow from MA mice and treated whole bone marrow cells *ex vivo* with RvD2 (1 nM) followed by methocult assay to examine myeloid potential. Colony growth was analyzed after 1 week of culture and myeloid colonies were enumerated (**Fig. 5F**). We observed a significant increase in macrophage colony forming units (CFU-M) whereas no changes were observed in granulocyte colony forming units CFU-G or CFU-GM (**Fig. 5G**), though a significant decrease in CFU-GEMM was seen. Therefore, the increase in MPP_GM_ *in vivo* and the striking increase in CFU-M *in vitro* indicate that RvD2 increased monocyte and macrophage potential in MA mice. Our data suggest that RvD2 acts directly on bone marrow cells to enhance monopoiesis and expedite differentiation of CFU-GEMM toward the monocyte lineage. Collectively, these data reveal that RvD2 treatment results in a systemic increase in monocytes that correlates with decreased steatosis and hepatic fibrosis in the liver, suggesting that RvD2 may protect the liver by reprogramming hematopoietic function.

### RvD2 limits age-related collagen accumulation in the liver in part through actions on bone marrow

To directly test the potential role of bone marrow function in driving the aging phenotype of the liver, and to test whether RvD2 could mitigate fibrosis via actions on bone marrow, we established bone marrow chimeric mice. Young recipient mice (CD45.1), aged 8-12 weeks, were lethally irradiated and reconstituted with congenic donor bone marrow from young (∼2 months) or old (17 months) mice, in combination with competitor donor bone marrow from young mice (**Fig. 6A**). To determine whether RvD2 could mitigate the impacts of old bone marrow, we treated donor bone marrow *ex vivo* with RvD2 (as above) and administered RvD2 to transplant recipients for one week following transplantation. Vehicle-treated controls received the same numbers of injections during the week-long treatment. All mice were allowed to equilibrate for four months where they received no further injections of RvD2 or Veh prior to sacrifice and analysis of liver collagen.

**Fig. 6.**
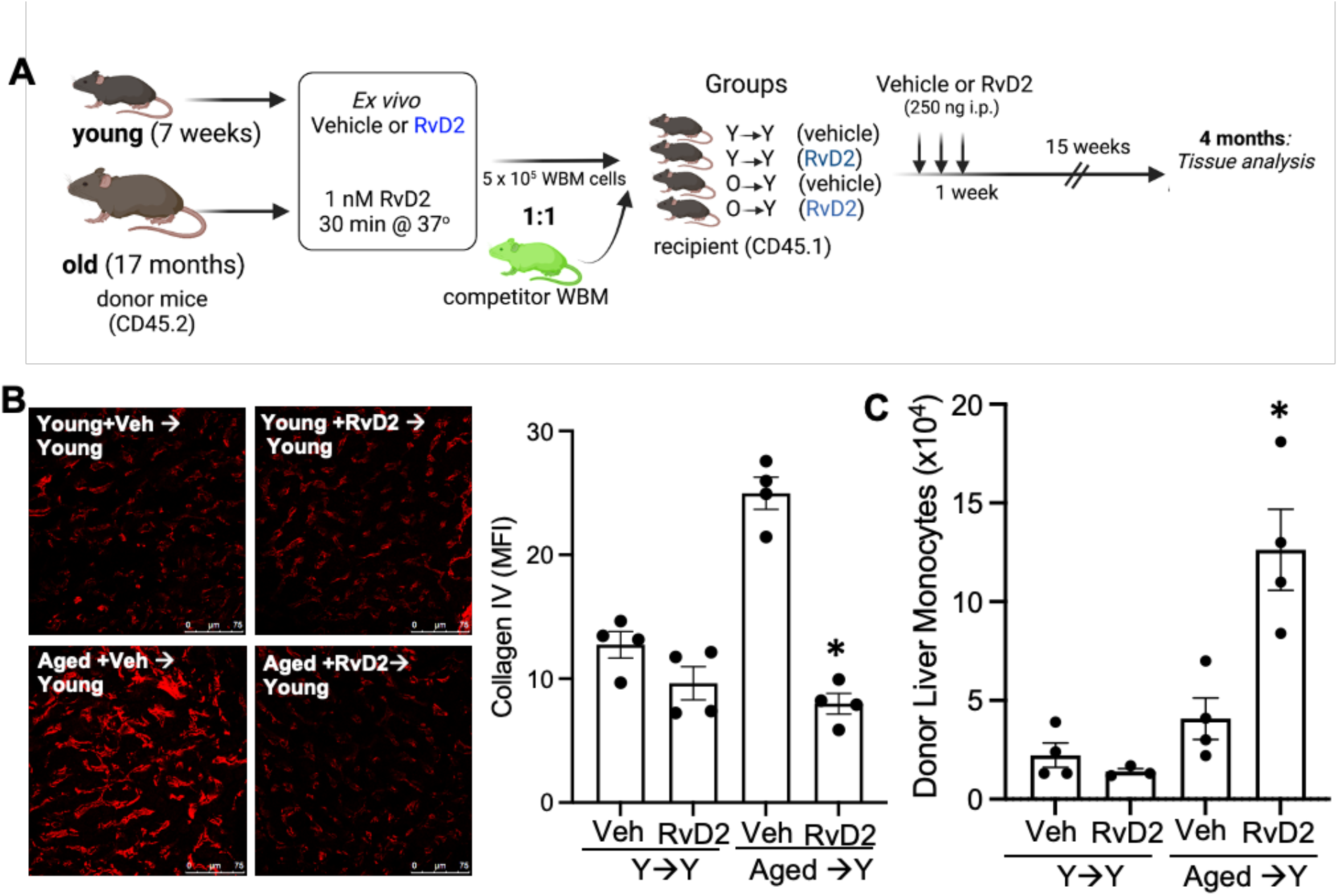
RvD2 limits liver age-Related collagen accumulation in part via changes in the bone marrow. (**A**) Schematic of the experimental set-up of competitive bone marrow transplants where young mice received either young or old donor bone marrow with or without RvD2 treatment. Recipient mice received injections of either vehicle or RvD2 for 1 week, and mice were analyzed 4 months post-transplant. (**B**) Representative images and quantification of liver sections stained with anti-Col-IV antibody shown in red. Images were acquired on a Leica confocal microscope and quantification of Col-IV MFI was done with Image J software. (**C**) Flow cytometry of liver samples was performed to identify donor-derived monocytes. Results are mean ± SEM and each dot represents an individual mouse. *p< 0.05, One way ANOVA with Kruskal-Wallis and a Dunn’s post-test.

In support of the idea that bone marrow-derived cells participate in age-associated liver collagen deposition we found that mice receiving transplants of old bone marrow exhibited pronounced Col-IV accumulation (**Fig. 6B**), relative to recipients of young bone marrow. However, RvD2 treatment had a marked impact on liver collagen in the recipients of old bone marrow, and we noted a significant reduction in Col-IV staining in this group, resulting in similar staining as recipients of young marrow (**Fig. 6B**). Moreover, we observed a significant increase in donor-derived monocytes in the livers of mice that received bone marrow from old donors and treated with RvD2 (**Fig. 6C**). Therefore, treatment with RvD2 was able to reverse the fibrotic phenotype in recipients of old bone marrow. These data demonstrate that a) the age of the donor bone marrow contributed to amount of collagen in the liver, and b) that even transient treatment with RvD2 was able to reverse this phenotype, correlating with increased monocytes.

## Discussion

Here, we report that *Gpr18* expression is elevated in macrophages from old mice and livers from elderly humans, suggesting a link between increased *Gpr18* expression and inflammaging. Myeloid-specific RvD2-GPR18 signaling limited age-related steatosis and hepatic fibrosis. Mechanistically, RvD2 treatment of MA mice resulted in a systemic increase in monocytes in the bone marrow, blood and liver that correlated with decreased steatosis and hepatic fibrosis, suggesting that RvD2 may protect the liver by reprogramming hematopoietic function.

Aging is associated with increased chronic low levels of pro-inflammatory factors(14). We found that *Gpr18* expression was increased in macrophages from old mice, which was associated with increased gene sets for “inflammation”, “map kinase signaling”, and “ROS”, as example. Pro-inflammatory ligands like toll-like receptor 4 agonists (e.g., lipopolysaccharide; LPS) and interferon-Ψ (IFNΨ) increase *Gpr18* mRNA expression in murine macrophages (26), which suggests that inflammation increases *Gpr18* expression. Conceptually, pro-inflammatory ligands like PGE_2_ are known to program resolution responses (27) and our results suggest that another feed-back mechanism may include a complex of pro-inflammatory ligands and pro-resolving GPCRs. How *Gpr18* expression is regulated is not known, but our work in the context of others suggest that pro-inflammatory factors and aging may drive its increased expression.

The actions and mechanisms of RvD2 in aging were not known prior to this study. However, Resolvin D1 (RvD1), which is another member of the D-series resolvin family, was shown to limit neutrophil infiltration into the peritoneum (2) and reduced neutrophils in the lungs after remote organ ischemia reperfusion injury (4) in old animals. Another study demonstrated that RvD1 treatment to aged mice reduced skeletal muscle inflammation and fibrosis, again suggesting a role for D-Series resolvins in limiting inflammation and tissue injury in aging (28). The mechanisms of actions of RvD1 in aging are not fully known. RvD1 binds and signals through distinct GPCRs called ALX/FPR2 (in mice and humans) and GPR32 (in humans) and a deeper exploration of these signaling axes in aging may yield important clue as to how neutrophils and other inflammatory factors may be regulated.

Our study focused on an RvD2-GPR18 signaling axis and we found that RvD2 treatment limited steatosis and hepatic fibrosis that was associated with increased monocytes in the bone marrow, blood and liver. Similarly, the SPM Maresin 2 was shown to increase liver monocytes in diet-induced obesity (DIO) young mice (29), which was associated with reduced inflammation. Whether monocyte recruitment to the liver in the context of obesity or aging is a common function for SPMs remains to be explored. Monocytes are highly dynamic in their functions and can promote damage as well as repair. In acute liver injury in young mice, a subset of monocytes transition into a reparative phenotype that aid in liver wound repair (30). Moreover, recruited monocytes can promote fibrosis resolution in mice, in part through the ability to promote phagocytosis and via expression of key matrix metalloproteinases (MMPs), like MMP9 (31). Therefore, stimulating a phagocytic macrophage phenotype from monocyte-derived macrophages protects against liver fibrosis. Indeed, there is ample evidence suggesting a predominant role for macrophages in the progression of liver fibrosis or repair, which is contextual and may hinge on the balance between tissue destructive and tissues reparative MMPs (18). How exactly RvD2 promotes the resolution of fibrosis is not known, but our data suggest that myeloid RvD2-GPR18 signaling is critical.

A recent study demonstrated that monocytes retained an inflammatory signature in the bone marrow during NAFLD progression (32). These findings suggest that bone marrow monocytes can be programmed to retain phenotypic information when in target organs. These data in the context of ours suggest that RvD2 may imprint bone marrow to exert a pro-resolving function in the liver. From an endogenous perspective, RvD2 is produced within the bone marrow upon ischemic inflammation (9) and so local production and signaling within the bone may be an additional mechanism for RvD2’s protective actions. Indeed, we found that RvD2 acted directly on the bone marrow to functionally produce monocyte progenitors in middle-age.

Indeed, aging is characterized by changes to blood production and immune responses that contribute to reduced vaccine efficacy, increased inflammation, and weakened host defenses seen in the elderly (33). Well-described changes to hematopoiesis include cell production that favors myeloid cell output including greater myeloid and megakaryocyte potential (33), though stem cell function declines. The mechanisms driving these changes have been linked to both microenvironmental signals from the bone marrow niche (34) as well as cell-intrinsic molecular changes (35-37). We, and others, have found that signs of hematopoietic aging are evident in middle age (38), which suggests MA may be an appropriate time for intervention. Furthermore, the ability of old bone marrow to promote a fibrotic phenotype in livers from transplants in young mice demonstrates the important contribution of hematopoietic cells in age-induced hepatic fibrosis. Importantly, the transient treatment with RvD2 in the transplant experiments demonstrated the durability of its effects, as hepatic fibrosis was significantly reduced more than three months after RvD2 treatment.

A major pro-resolving feature of RvD2 in our context was reduced steatosis and hepatic fibrosis in MA mice. SPMs limit several features of liver disease in young mice (reviewed here (39)). Some notable examples include that LXA_4_ limits diet-induced liver triglyceride, ALT levels (40) suggesting liver protective role in young mice fed a high fat diet. Also, resolvin D1 (RvD1) limits steatosis and fibrosis in the MCD NASH model (41). Therefore, SPMs may be an intriguing therapeutic strategy in the context of liver diseases.

An important problem with anti-inflammatory treatment strategies is that they reduce host defense mechanisms. Indeed, blocking inflammation can lead to increased susceptibility of infections, which is especially critical in the elderly population, and so new ways to control inflammation without compromising host defense is an urgent clinical need. The RvD2-GPR18 signaling axis has been shown to boost host defense in mice. Indeed, evidence for the RvD2:GPR18 interaction *in vivo* came from studies of murine self-limited inflammation initiated by zymosan or *Escherichia coli* (*E. coli*). In mice with global deficiency of *Gpr18*, neutrophil infiltration during bacterial peritonitis is higher than that of WT littermate controls, leading to delayed resolution of acute inflammation (8). This is associated with a defect in *E. coli* phagocytosis as well as macrophage efferocytosis. Administration of RvD2 expedites resolution and enhances efferocytosis when given at the peak of inflammation. These actions are not observed in GPR18-knockout (KO) mice, indicating that these resolution-enhancing roles of RvD2 are GPR18-dependent *in vivo*. Similarly, GPR18 is required for RvD2-dependent resolution of inflammation and enhancement of host-defense in a distinct model of skin infection initiated by *Staphylococcus aureus* (8). Overall, our work suggests a new function for RvD2 in aging and may prove to be an intriguing therapeutic in the context of age-related liver disease.

## Material and Methods

### Experimental animals

All animal experiments were conducted in accordance with the Albany Medical College IACUC guidelines for animal care and were approved by the Animal Research Facility at Albany Medical College. Male C57BL/6 young mice (2-3 month), middle-aged mice (11-12 months), old (17-19 months), and UBC-EGFP mice were purchased from The Jackson Laboratory. Male C57BL/6.SJL (expressing the *Ptprc* a allele; CD45.1) were purchased from Taconic Biosciences. Mice were housed in the Albany Medical College Animal Research Facility and fed a standard rodent chow diet during the experiment. Mice were socially housed in standard cages at 22°C under a 12 hr light and 12 hr dark cycle.

### Generation of a humanized *GPR18* floxed mouse

Murine *Gpr18* was removed and replaced with human *GPR18* that was flanked with two loxp sites. The original mice contained a NeoSTOP cassette, which was removed upon breeding with a FLPe mouse flippase (B6.129S4-*Gt(ROSA)26Sortm1(FLP1)Dym*/RainJ; Stock Number: 009086, JAX). Once the NeoSTOP was successfully removed, these mice were then backcrossed with C57BL6 mice. After successive backcrossing with C57BL6 mice, human *GPR18* was confirmed with genotyping and mRNA expression via qPCR. These human GPR18 floxed mice, are referred to in the text as “fl/fl” mice. These fl/fl mice were then crossed with a lysozyme Cre (LysM, JAX stock #004781) to remove human GPR18 in myeloid cells. Successive rounds of breeding occurred to ensure the loss of murine Gpr18 and the loss of human GPR18 on myeloid cells, referred to in the text as mKO (or myeloid knockout).

### RvD2 treatment

11-month-old male C57BL/6 mice were purchased from The Jackson Laboratory. Mice were intraperitoneally (i.p.) injected with either vehicle (PBS) or 250 ng of RvD2 (Cayman Chemicals Cat#10007279), for 7 days consecutively. The mice were sacrificed on day 8, and their body weight and complete blood count were measured, peritoneal cells were isolated by lavage, and liver was collected for end-point analysis as described below.

### Complete Blood Count

Whole blood from mice was collected in 10% EDTA tubes. Complete blood count was measured immediately using a Heska-Element HT5 hematology analyzer.

### RNA sequencing

Male C57BL/6 young (age 2 months) mice or old (18 months) mice were purchased from The Jackson Laboratory. Mice were then i.p. injected with 200 µg of ZymA (or ZymA, Sigma, Cat #Z4250) per mouse. After 72 hours, peritoneal cells were harvested by lavage and cultured overnight in complete DMEM containing L-cell media. The following day, unattached cells were washed with PBS mRNA was extracted from the attached macrophages with a Qiagen RNeasy Mini kit (Cat #74106) according to the manufacturer’s instructions. RNA sequencing was performed at The Forsyth Institute. There were 3 mice in each group and RNA from each mouse was sequenced in triplicate. The gene counts were normalized by rlog transformation and analyzed by the DeSeq2 pipeline. The analyzed DeSeq2 results were uploaded on the web-based application called iPathway Guide and significantly upregulated and downregulated genes and pathways were identified. The log2fold change threshold was >0.6 and the p-value threshold was <0.05 was applied. Moreover, the Gene Set Enrichment Analysis (GSEA) was performed using the default settings.

Bone marrow and liver RNA sequencing: The microarray data of 23 aging human male and female liver, were downloaded from the Gene Expression Omnibus (GEO, https://www.ncbi.nlm.nih.gov/geo/) database, accession number GSE133815. In addition, the GSE98249 data set was downloaded with bulk RNA-seq data of 6 young and 6 aging CD11b+ macrophages from murine bone marrow. The expression level of each gene was transformed into a log2 base before further analysis. Differential gene expression was analyzed using DESeq2 (version: 3.16) (42), in R studio software. Significance was defined as p-value < 0.05.

### Liver Picrosirous Red Staining

Livers were perfused and isolated from each mouse, fixed in 4% paraformaldehyde (PFA), and embedded in OCT. 10μm thick liver sections were cut with a Leica Biosystems cryostat and mounted on glass slides. Liver sections were gradually dehydrated in xylene and ethanol and were stained by hematoxylin and eosin staining. Further, collagen content in the liver was determined by picrosirius red staining as per the manufacturer’s instructions (Polysciences, Cat#24901). After staining, sections were mounted with aqueous mounting media and were examined under the microscope (Olympus DP74). Alterations in histology between the three groups were assessed and quantified using Image J.

### Liver Immunofluorescence

Murine livers were perfused, isolated and placed in 500 μL of 4% PFA overnight at room temperature. The livers were then placed in 500 μL of 30% sucrose solution for 48 hours at 4°C, then embedded in OCT. Murine livers were sectioned into 10-12 μm cross sections using a Leica Cryostat Machine (CM1860) at and stored at -80°C. Liver sections were at stained at 4°C with anti-Collagen IV (EMD Millipore Cat# 3607063) and anti-Galectin 3 (Mac-2) (Cedarlane Cat# CL8942AP). For this co-stain, an antigen retrieval method was added for optimal antibody binding and visualization. Briefly, frozen sections were first placed in a slide holder and under a ventilation hood to dry for 15 min. The sections were then submerged in 1x PBS to rehydrate for at least 5 min, followed by a conventional rehydration using sequential incubations with xylene, 100%, 95%, 70% ethanol, 50% and distilled H_2_0 each for 1 min. The slides were then submerged into Antigen Retrieval Buffer (Antigen Unmasking Solution Citric Acid Based, # H-3300, Vector Biolabs) and the entire unit was placed in a 100°C water bath 15 min. The unit was then removed from the hot water bath and left at room temperature for 30-min cooling period. The sections were blocked with 1% BSA and 10% goat serum for 1h at 4°C, followed by washes in 1x TBST 0.1% TWEEN buffer 3 times for 5 min each. Primary antibodies were added to the slides in 1% BSA overnight at 4°C. After 24h, slides were washed with 1x TBST 0.1% TWEEN buffer 3 times for 5 mins each. Secondary antibodies were added in 1% BSA for 2h at 4°C, followed by a PBS wash. Finally, the sections were counterstained with Hoechst to identify nuclei. The slides were visualized on a Leica SPE confocal microscope. The mean fluorescence intensity and cell count per visual field was determined through analysis using ImageJ software.

For the transplantation experiments, murine livers were isolated and flash frozen in liquid nitrogen and then stored at -80°C. These livers were then sectioned into 10-12 μm cross sections using a Leica Cryostat Machine (CM1860) at and stored at -20°C. Liver sections were stained at 4°C with anti-Collagen IV (EMD Millipore Cat# 3607063). Frozen sections were submerged in ice-cold 100% Methanol for 10 min. The sections were washed with PBS and then immediately blocked with 1% BSA and 10% goat serum for 1h at 4°C. The sections were washed with PBS to remove the blocking reagent and primary antibodies were added in 1% BSA overnight at 4°C. After 24 h, slides were washed with PBS and secondary antibodies were added in 1% BSA for 2 h at 4°C. The sections were washed again and counterstained with Hoechst to identify nuclei. The slides were visualized on a Leica SPE confocal microscope.

### Oil Red O staining

Liver sections were fixed in 4% PFA for 5 min and then washed with 60% isopropanol for a further 5 min. Next, liver sections were stained with freshly prepared Oil red O (Sigma, Cat#O0625) staining solution for 30 min. Following incubation, they were washed with 60% isopropanol for 2 mins and immediately rinsed in distilled water. Stained liver sections were mounted with Immumount and images were viewed on 20x objective lens using an Olympus DP74 microscope and Olympus DP2-BSW software. Oil Red O positive areas were quantified by threshold analysis using ImageJ.

### Triglyceride assay

Livers were perfused, isolated and immediately snap frozen in liquid nitrogen. Liver homogenates were subjected to triglyceride assay (Cell Biolabs, cat# STA-396) according to the manufacturer’s instructions.

### Tissue Processing and Flow Cytometry

Perfused livers were isolated and processed by mincing with scissors, followed by enzymatic digestion. Liver mononuclear cells were isolated using the previously reported method of *in vitro* 0.05% collagenase incubation of liver specimens (43-45). Briefly, the liver was perfused with 10 mL ice cold PBS via the portal vein. Immediately after perfusion, the liver was removed and minced with scissors. Minced livers were centrifuged, and supernatant was removed. Homogenized livers were resuspended in 10mL HBSS containing 0.05% collagenase (Type IV; Stem cell technology; cat#07427) and the specimens were shaken for 20 min at 37°C. Digested liver tissue was centrifuged at 450g for 5 min at 4°C, and liver tissue was filtered through a 70uM filter. Filtered liver tissue was resuspended in 10 mL 33% Percoll solution (Sigma-Aldrich; Cat#P1644) and centrifuged at 600g for 20 min at room temperature (with brake set to 0). The supernatant was aspirated, and cell pellets were lysed with 200uL of ACK buffer for 4 min at RT. After RBC lysis buffer, cells were washed twice and resuspended in HBSS containing FBS and counted. Single cell suspensions were plated and stained after incubation with Fc block using antibodies directed against myeloid cell markers (CD45.2 PerCP Cy5.5; F4/80 APC; Ly6C BV510; Ly6G BV605; CD115 PE; CD11b PE Cy7; Supplemental Table S1). Data were acquired on a FACSymphony flow cytometer (BD Biosciences) or a Northern Lights spectral flow cytometer (Cytek Biosciences) and analyzed using FlowJo (TreeStar) software.

Bone marrow was flushed from both hindlimbs and filtered through a 70 um mesh filter. After red blood cell lysis (ACK lysis buffer), single cell suspensions were plated and stained for flow cytometry. Cells were Fc blocked and then stained for HSPCs using antibodies against lineage markers (CD3, CD11b, Gr-1, Ter-119, CD45R), and HSPC markers (c-Kit, Sca-1, CD150, CD48, and CD135; Supplemental Table S1). Cells were also stained with markers against myeloid cells (CD11b, Ly6C, Ly6G, CD115, F4/80, CD115; Supplemental Table S1).

### Colony Forming Unit Assays

Bone marrow single cell suspensions were first incubated for 30 minutes at 37°C in vehicle (define) or RvD2. Cells were washed with 1x PBS and plated in methocullulose media (Methocult M3434, Stem Cell Technologies). Cells were plated (4 × 10^5^ cells per 35 mm-tissue culture dish) in duplicate, and plates were incubated for 7-8 days at 37°C in 5% CO_2_. Plates were evaluated and myeloid colonies were distinguished and counted under a light microscope.

### Bone Marrow Transplantation Experiments

Recipient 10-week-old C57BL/6.SJL male mice were treated with antibiotics (sulfamethoxazole and trimethoprim diluted in drinking water (200mg/40mg in 5ml) for 10 days) prior to transplantation. Recipient mice were lethally irradiated with a total of 950 rads (split-dose; 475 rads, 24 hours apart) and then reconstituted with a total of 5 × 10^5^ total bone marrow cells. Mice were maintained on antibiotics for 15 days post-transplantation and then placed on normal drinking water. Donor bone marrow was a 1:1 mixture of young whole bone marrow from UBC-EGFP mice and experimental bone marrow from donor young (7-week-old) or old (18-month-old) bone marrow either treated with vehicle (PBS) or RvD2. Donor bone marrow from young or old mice was incubated at 37°C for 30 minutes in vehicle or 1 nM RvD2 then washed with 1x PBS prior to transplant. Transplant recipients then received intraperitoneal injections of either vehicle (PBS) or RvD2 (250 ng/mouse/day) for 5 days. Mice were allowed to recover and analyzed 4 months post-transplantation. See schematic in Figure 6A. Livers were harvested from transplant recipients and analyzed to distinguish donor cells and myeloid populations (as described above).

### Statistical analysis

MA mice were randomly assigned to receive either vehicle or RvD2 groups. Statistical differences between the *in vivo* groups were determined using either the two-tailed Student’s t-test, Mann-Whitney or Kruskal Wallis with a Dunn’s post-test analysis using GraphPad Prism. All results are represented as mean ± S.E.M. A p-value equal to or lower than 0.05 was considered statistically significant. The figure legends have details regarding respective statistical tests.

## Authors Contributions

G.F., H.F., J.B., S.S., H.A.M, M.M. performed *in vivo* experiments a. S.S. performed and analyzed the bulk RNAseq from young vs aged macrophages. R.B. and A.A. performed informatics analysis on human livers. H.F. and C.D. conducted and analyzed picrosirious red and Oil red O staining of liver. M.L. and H.F. conducted and analyzed the TG assays. H.F., M.N., K.G. conducted and analyzed liver immunofluorescence and ELISA analyses. J.B. and N.B. conducted bone marrow and liver flow cytometry, and J.B. performed methocult assays, and performed and analyzed transplant experiments. M.S. supervised selected experiments and contributed to the writing of the manuscript. G.F. K.C.M., H.F., J.B. contributed to the writing of the manuscript and K.C.M. and G.F. conceived the overall design of the experiments. The order of the first authors was determined on the basis of their effort and contributions to the study.

