## Supplemental Methods and Data for "The Resolvin D2-GPR18 Axis Enhances Bone Marrow Function and Limits Hepatic Fibrosis in Aging"

### Supplementary Information

### Methods:

#### Immunofluorescence (*in vitro*)

Peritoneal Macrophage staining - On the day of sacrifice, peritoneal cells were isolated from either young vehicle, old vehicle, or old RvD2 mice by lavage, and cultured overnight in complete DMEM containing L-cell media for macrophage differentiation. Peritoneal macrophages were fixed with 4% PFA for 15 mins, washed with 1X PBS, and permeabilized with 0.1% Titron-X-100 for 15 mins at room temperature. Fixed cells were blocked with blocking buffer (5% BSA in PBS + 0.1% Titron-X-100 + 5% goat serum) and stained with rabbit anti-mouse p-p38 primary antibody at 1:200 dilution (Cell signaling, cat #9211), yH2AX (Cell Signaling, 9211S) and COX2 (Cell Signaling, 12282S) overnight at 4°C. The next day, cells were washed with 1X PBS and incubated with goat anti-rabbit Alexa Fluor-594 secondary antibody (Invitrogen A-11037) at 1:250 dilution for 2 hours at RT. Cell nuclei were stained with Hoechst (1ug/ml) for 10 mins and images were acquired immediately on a Leica SPE confocal microscope. 6-7 different fields were acquired per well per antibody per mouse.

#### CRP ELISA

Plasma was collected by spinning whole blood for 30 mins, 21,000 rcf at 4°C. Plasma was assessed for CRP by ELISA analysis according to the manufacturer (Thermo Fisher, #EM20RB).

### Antibody List

| Marker | Species | Fluorophore | Clone | Company (Cat. No.) |
|---|---|---|---|---|
| F4/80 | | APC | CI:A3-1 | Abcam (AB105080) |
| Ly6C | | Pacific Blue | HK1.5 | BioLegend (128013) |
| Ly6G | | APC-Cy7 | 1A8 | BioLegend (127623) |
| CD11b | | PE-Cy7 | M1/70 | BioLegend (101216) |
| Gr-1 | | FITC | RB6-8C5 | BioLegend (108406) |
| CD11b | | FITC | M1/70 | BioLegend (101206) |
| B220 | | FITC | RA3-6B2 | BioLegend (103206) |
| Ter-119 | | FITC | TER-119 | BioLegend (116206) |
| CD3 | | FITC | 17A2 | BioLegend (100204) |
| c-Kit | | APC | 2B8 | BioLegend (105812) |
| CD48 | | Pacific Blue | HM48-1 | BioLegend (103418) |
| CD41 | | BV510 | MWReg30 | BioLegend (133923) |
| CD150 | | BV711 | TC15-12F1 | BioLegend (115941) |
| CD135 | | PE | AF210 | BioLegend (135306) |
| Sca-1 | | PE-Cy7 | D7 | BioLegend (108114) |
| CD115 | | PE | AFS98 | BioLegend (135506) |
| CD45.2 | | BV711 | 104 | BD Biosciences (563685) |
| Collagen IV | Rabbit | | ATS2 | EMD Millipore (AB756P) |
| Mac2 | Rat | | M3/38 | Cedarlane Labs (8942AP) |
| COX2 | Rabbit | | D5H5 | Cell Signaling (12282S) |
| p-p38 | Rabbit | | T180/Y182 | Cell Signaling (9211S) |
| gH2AX | Rabbit | | S139 | Cell Signaling (2577S) |

### Supplementary Figures

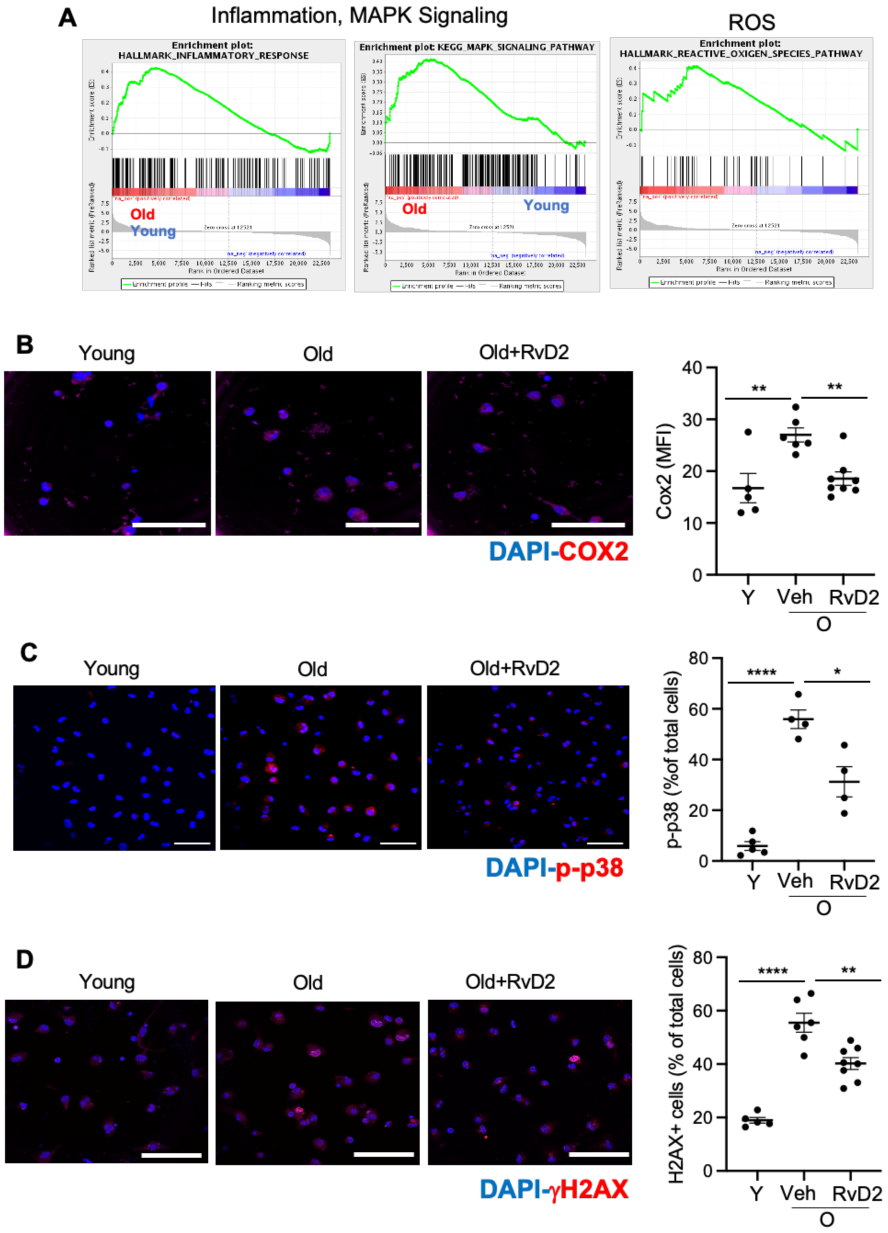


**Supp Fig 1. Macrophages from aged mice have an increased COX2 p-p38 and γH2AX and decreased by RvD2.** (**A**) GSEA analysis from the RNA sequencing performed in Fig. 1 is shown. Peritoneal macrophages from young, old or, old mice treated with RvD2 (described in the methods) isolated, cultured overnight, and immuno-stained for (**B**) COX-2, (**C**) p-p38 or (**D**) H2AγX are shown in magenta, whereas nuclei are stained in blue with DAPI. Magnification is 40X and the scale bar is 50μm. ****p<0.0001, ns-non-significant. One-way ANOVA with Tukey's multiple comparison test. Each symbol represents an individual mouse. Results are mean ± S.E.M.

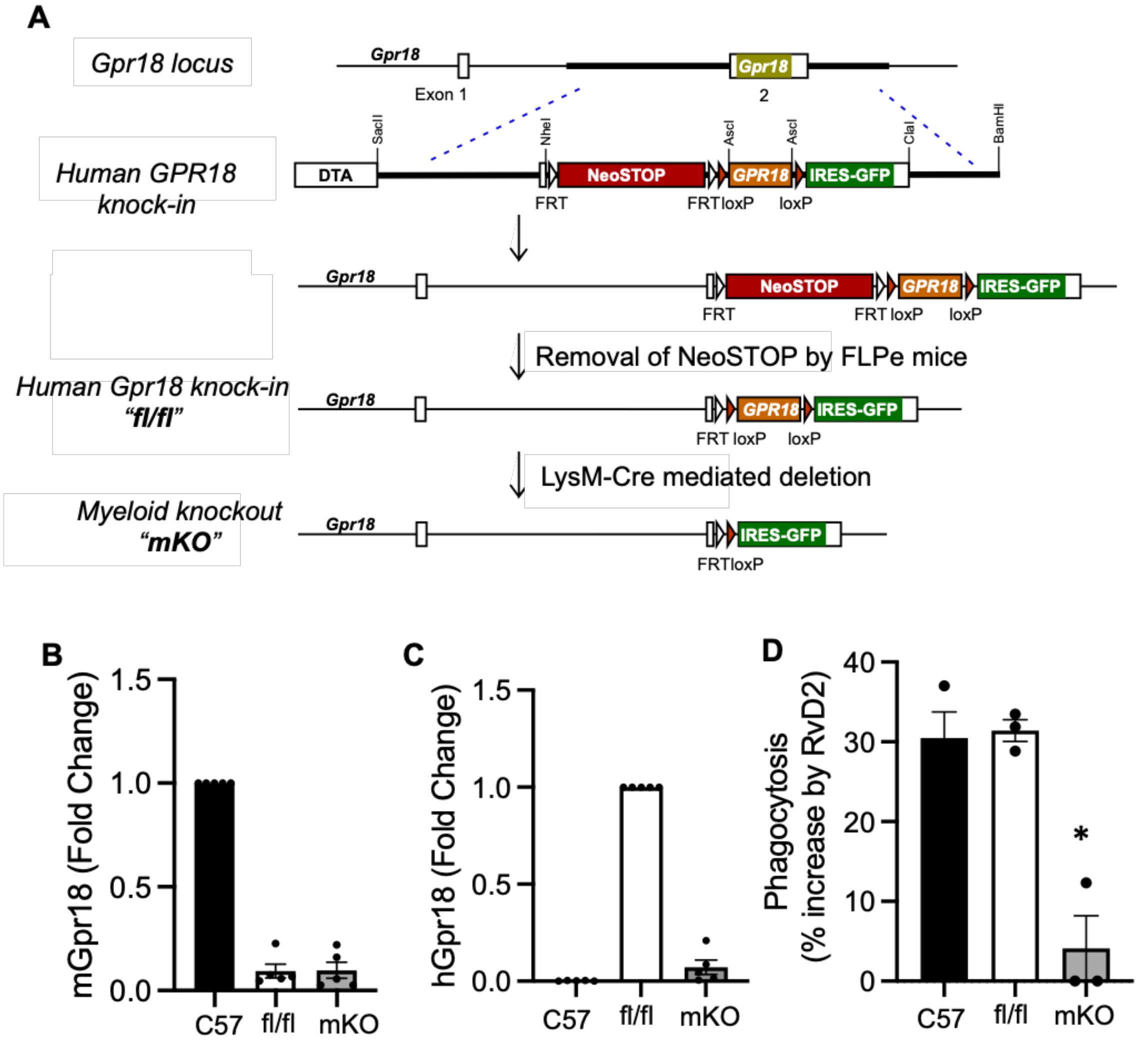


**Supp Fig 2. Generation of humanized GPR18 floxed mice.** (**A**) Scheme of humanized Gpr18 floxed and Gpr18-Lyzm (mKO) mice were made as described in the methods section. (**B-D**) BMDMs from c57Bl7/6 mice, fl/fl and mKO mice were subjected to qPCR and assessed for levels of (**B**) murine *Gpr18* and (**C**) human *GPR18*. (**D**) Phagocytosis was carried out as described in the methods and data were assessed by fluorescence plate reader and represented as percent increase of phagocytosis by RvD2.

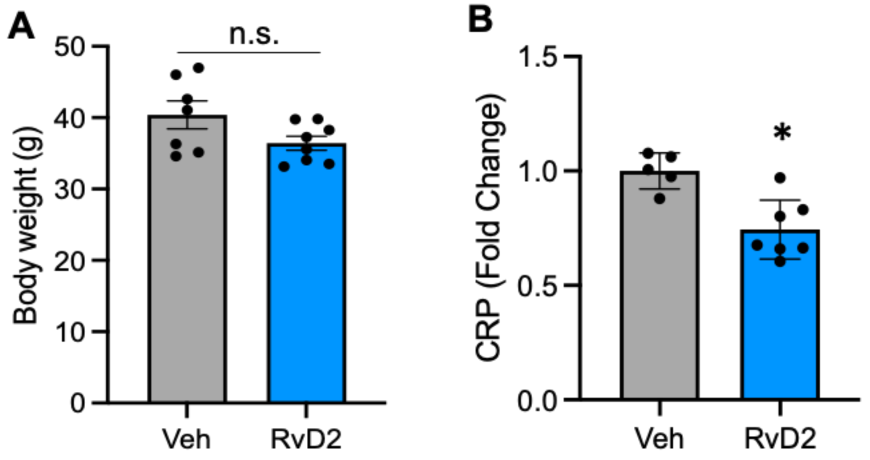


**Supplemental Fig 3. RvD2 treatment to MA mice does not change body weight and limits CRP.** (**A**) MA mice were treated with Veh or RvD2 as described in the methods and body weight were calculated. (**B**) Plasma was collected and subjected to ELISA analysis for circulating levels of CRP.

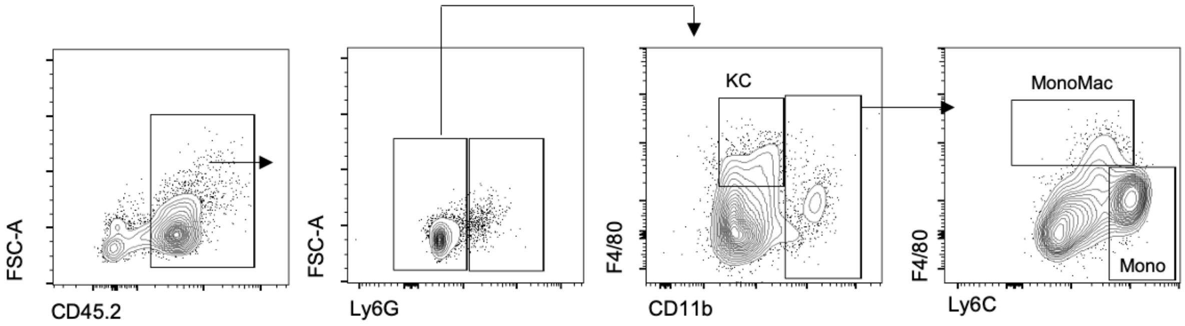


**Supp Fig 4. Gating Strategy for liver myeloid cells.** Livers were processed for flow cytometry and stained with antibodies against myeloid markers, followed by analysis. Kupffer cells were identified as CD45.2$^{+}$, Ly6G$^{-}$, CD11b$^{lo/-}$, F4/80$^{hi}$; Monocyte-Macrophages were identified as CD45.2$^{+}$, Ly6G$^{-}$, CD11b$^{+}$, Ly6C$^{lo/-}$, F4/80$^{+}$; Monocytes were identified as CD45.2$^{+}$, Ly6G$^{-}$, CD11b$^{+}$, Ly6C$^{hi}$. Cells were analyzed on a BD FACSymphony or a Cytek Northern Lights instrument.

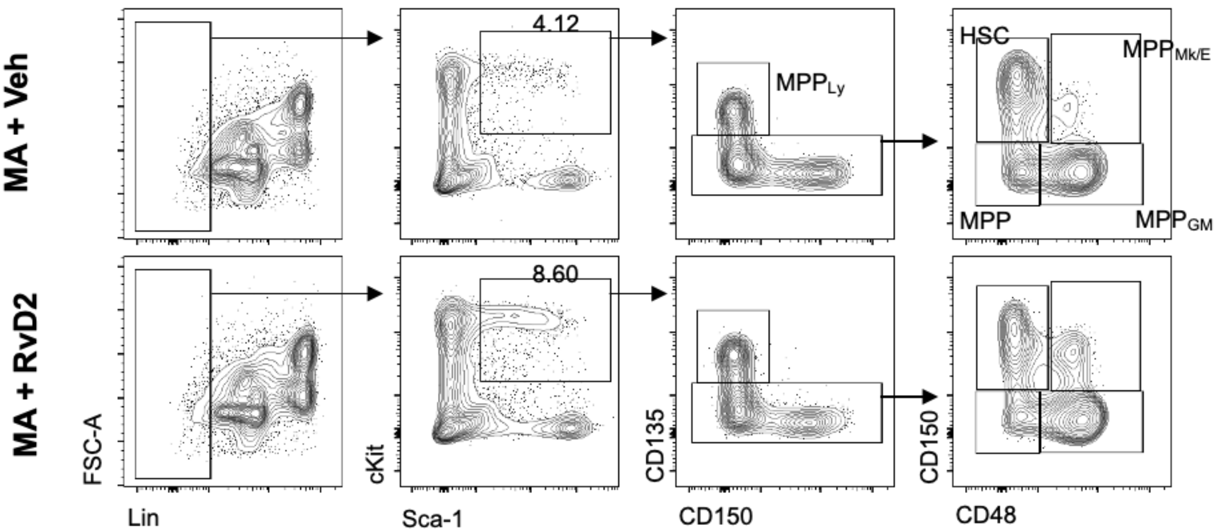


**Supp Fig 5. Gating strategy for bone marrow HSPCs.** Bone marrow was isolated from the hindlimbs of MA + Veh and MA + RvD2 mice. Single cells were first gated on lack of lineage markers (CD45R, Gr-1, Ter-119, CD11b, CD3). HSPCs were identified as Lin-, cKit+, Sca-1+ (LSK). $MPP_{Ly}$ cells were identified as $CD150^-$, $CD135^+$. HSCs were $CD135^-$, $CD150^+$, $CD48^-$; $MPP_{Mk/E}$ cells were $CD135^-$, $CD150^+$, $CD48^+$; $MPP_{GM}$ cells were $CD135^-$, CD150-, $CD48^+$; MPPs were $CD135^-$, $CD150^-$, $CD48^-$. Cells were analyzed on a BD FACSymphony or Cytek Northern Lights instrument.
